# Genetic Modeling of Dyadic Behavioral Traits: Implications for Estimation and Interpretation of Variance Components

**DOI:** 10.64898/2026.06.10.731434

**Authors:** Xiaohan Jiang, Janice M Siegford, Juan P Steibel

## Abstract

Studying the genomic control of dyadic social interactions is gaining traction in animal genetics. However, genetic modeling of social interactions poses several challenges, one of which is whether social interactions should be treated as dyadic traits or as aggregated traits at the individual level. In this study, we systematically compared two approaches: dyadic models using dyadic traits and marginal models using marginally aggregated traits and we derived the algebraic relationships between their variance components. In the application, we used a published dataset on post-mixing aggression in pigs, including both directed and undirected aggression records collected during the 9-hour period after mixing among 797 finishing pigs in 59 social groups, as an example to show how model choice can affect variance estimation. Results showed that dyadic models can estimate genetic effects and permanent environmental effects by exploiting repeated dyadic interaction records, thereby enabling a more complete understanding of the sources of variation underlying social interactions. In contrast, marginal models can bias the estimation and interpretation of genetic components, as the aggregated genetic variance may be confounded with other variance components due to the aggregation of dyadic traits. Marginal models may also lead to overestimation of social group and residual variance. These results can provide useful guidance for choosing appropriate modeling strategies for social interaction traits.

## Introduction

Most farm animals live in groups and engage in various forms of social interaction, which play a crucial role in shaping their performance, health, and welfare. There is substantial evidence that genes can influence complex social behaviors. For example, feather-pecking behavior in laying hens harms groupmates, while genetic selection against it can reduce it over generations (Kjaer et al. 2001). Thus, geneticists are interested in dissecting the components that affect social interactions to incorporate novel traits into breeding programs to select animals better adapted to group-housed production (Canario et al. 2020).

With the rapid advancement of sensing technologies for animal phenotyping, large-scale automated records of social interaction phenotypes are becoming available. For instance, in laying hens, Perinot et al. (2025) used spatial and temporal tracking data based on RFID systems to infer social associations in large aviaries. In dairy cattle, real-time location systems were used to continuously monitor dyadic spatial associations on dairy farms (Marina et al. 2024). Also, computer vision enables accurate tracking of animals’ movements and identification of their social behaviors (Hollifield et al. 2024; Agha et al. 2025). Hence, it is important to revisit models for dissecting genetic and environmental components of behavioral interactions.

The fundamental observational unit of social interactions is a pair of animals, which is called a “dyad” (Kenny et al. 2001). Thus, measuring social interactions yields dyadic phenotypic data. These dyadic data can be either directed or undirected. In directed dyadic data, two individuals play different roles: the giver (who actively delivers a behavior) and the receiver (the recipient of the behavior). Undirected or reciprocal interactions, in contrast, refer to the mutual involvement of the two individuals, with no specific giver or receiver identified. For example, in pig aggression, both interaction types are frequently observed (O’Malley et al. 2022).

A common approach to analyzing dyadic data is to sum two-sided interactions into individual-level observations (Han et al. 2022). For instance, in pig aggression, Turner et al. (2009) aggregated the total duration of aggression delivered and received by each individual pig by summing row-wise and column-wise the matrix of directed dyadic attacks. We call the resulting trait the marginal social trait and their model the marginal model. Moreover, the act of bullying, i.e.: summing over total aggression delivered, has been reported to be more heritable than the receipt of attacks (Løvendahl et al. 2005; Turner et al. 2009).

An alternative approach that is gaining momentum in the genetic analysis of dyadic interaction matrices is based on marginalization using social network analyses. For instance, Agha et al. (2025) analyzed AI generated proximity matrices as the adjacency matrix of a social network analysis and extracted individual (marginal) node statistics (e.g.: in- and out-degree, centrality, etc.) as social phenotypes and then analyzed them with classic animal genetic models (Agha et al. 2020).

All the cited approaches focus on modeling the genetic effects of marginal social traits, attributable to individuals rather than to a pair of animals, potentially leading to the loss of important information embedded in the dyadic nature of recorded social interactions. Moreover, it has been shown that social interactions are strongly influenced by their social environment and early-life experiences (D’Eath and Lawrence 2004; Canario et al. 2017). Such sources of persistent non-genetic variation may be captured statistically by permanent environmental terms. Thus, aggregating traits (marginal sums of dyadic interaction matrices) may omit important effects and lead to biased estimates of key variance components.

Quantitative frameworks for modeling dyadic social interactions have been developed. Løvendahl et al. (2005) analyzed the delivery and receipt of aggression of sows at mixing with a dyadic model. More recently, Han et al. (2022) presented a generalized linear mixed model (GLMM) framework to estimate phenotypic giver and receiver random effects for the duration of aggression. Finally, Wang et al. (2023) simulated two correlated traits corresponding to an individual’s tendency to give and to receive social interactions, including both genetic and permanent environmental effects.

Although dyadic models have been increasingly applied to social interaction data, their theoretical relationship with aggregated-trait models (marginal models) remains unclear. In addition, the formulation of directed and undirected dyadic models requires further clarification. Thus, the objectives of this study are: 1) to compare the theoretical similarities and differences between the dyadic and marginal social interaction models. 2) To illustrate the use of these models for the dissection of genetic and permanent environmental contributions in pig aggression.

### Theoretical Background

Assume that all interactions between animals are observed within multiple social groups. In the *k*^th^ group composed of *n* animals, dyadic interaction matrix ***Y***_***k***_ is defined as the matrix of pairwise interaction between every possible pair of individuals.

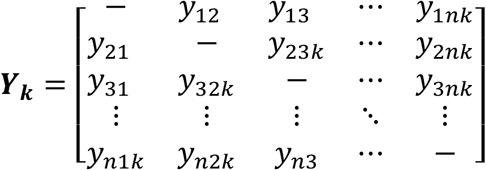

Directed interactions are interactions initiated by one animal toward another, with the behavior giver (rows of the matrix) and receiver (columns of the matrix) clearly distinguished. This type of interaction is commonly observed when a more dominant animal directs interactions toward a more submissive animal. Each animal, therefore, has two distinct traits: one for giving the interaction and one for receiving the interaction. For directed interactions, each off-diagonal element *y*_*ijk*_ represents the interaction phenotype from the giver *i* to the receiver *j*. Accordingly, ***Y***_***k***_ is asymmetric, with *y*_*ijk*_ ≠ *y*_*jik*_.

Undirected interactions, in contrast, are mutual interactions between two individuals, with no distinct giver and receiver roles. Each animal, therefore, has a single interaction trait. For example, social associations between animals obtained from location and tracking technologies can be treated as undirected interaction traits (Perinot et al. 2025). For these interactions, *y*_*ijk*_ represents the interaction phenotype between animal *i* and animal *j*. As a result, ***Y***_***k***_ is symmetric, with *y*_*ijk*_ = *y*_*jik*_.

In a group of size *n*, there are *n*(*n* − 1) possible directed dyads or *n*(*n* − 1)/2 possible undirected dyads, as self-interactions are not considered (diagonal elements *y*_*iik*_ are undefined). The interaction phenotype *y*_*ijk*_ could be binary outcomes (0/1 for no interaction/interaction), discrete outcomes (e.g., frequencies/counts of number of interactions), ordinal outcomes (none, low, medium, and high score of interaction on a Likert scale), or continuous outcomes (e.g., measured intensity or duration of interaction). The type of outcome will affect the distributional assumptions of the presented models. Without loss of generality, in this paper, we assume that the interaction is a continuous measure that can be modeled as a sum of Gaussian distributions (a linear Gaussian mixed model).

### Dyadic model for directed social interactions

In the dyadic social interaction model, direct dyadic social interactions can be partitioned into a giver’s additive genetic, a receiver’s additive genetic, a giver’s permanent environmental, and a receiver’s permanent environmental component. Thus, the dyadic genetic model can be interpreted as a social genetic effects (SGE) model (Bijma et al. 2007) with group size *n* = 2, while separately accounting for genetic effects (representing inherited components of behavior) and permanent environmental effects, which reflect prior experiences shaping current behavior. Unlike traditional SGE models, where the trait is typically an individual-level production trait representing the consequence of an animal acting as the receiver, the dyadic model analyzes pairwise behavioral interaction traits observed between two individuals, which is described as:

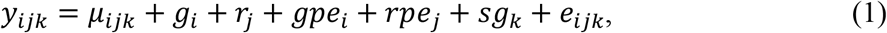

where *y*_*ijk*_ is the observed social interaction phenotype delivered by the *i*th animal (giver) to the *j*th animal (receiver) in the *k*th group. *µ*_*ijk*_ is the expected mean associated with *i* and *j. g*_*i*_ and *r*_*j*_ denote the random additive genetic effects of the giver *i* and the receiver *j*, respectively. *gpe*_*i*_ and *rpe*_*j*_ represent the random permanent environmental effects of the giver *i* and the receiver *j*, respectively. *sg*_*k*_ is the random effect of the social group *k*, assumed to be 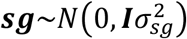, where 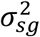 is the social group variance. *e* is the error, distributed as 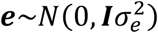, where 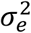 is the error variance. The ***g*** and **r, *gpe*** and ***rpe*** are assumed to be:

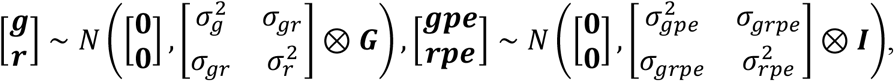

where ***G*** is the genomic or numerator relationship matrix, 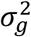 and 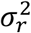 are the giver and receiver genetic variance, *σ*_*gr*_ represents the covariance between the giver and the receiver genetic effects of the same animal. ***I*** is the identity matrix, 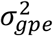 and 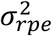 are the giver and the receiver permanent environmental variance, *σ*_*grpe*_ represents the covariance between the giver and the receiver permanent environmental effects of the same animal.

Based on Equation (1), the total variance of *y*_*ijk*_ is:

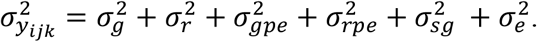

The heritability for giving interaction 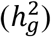 and receiving interaction 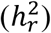 are defined as:

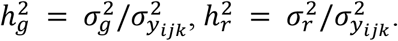

The repeatability for giving interaction (*r*_*g*_) and receiving interaction (*r*_*r*_) are expressed as:

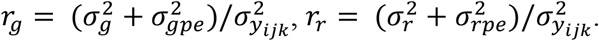

### Comparison with the marginal trait model

The Dyadic interaction matrix ***Y***_***k***_ can be summed row-wise or column-wise to get the individual-level total social interaction phenotype. The marginal *y*_*i*⋅*k*_ or *y*_⋅*jk*_ have been extensively used in previously published research (Turner et al. 2009). In this section, we study the implications of summing rows or columns of the behavior interaction matrix given different plausible assumptions for the involved variance components.

#### Row-Sum giver marginal Model

The giver marginal model is obtained by summing the dyadic interaction matrix ***Y***_***k***_ over rows:

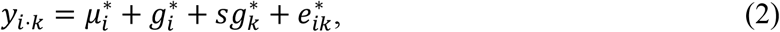

where *y*_*i*⋅*k*_ denotes the aggregated social interaction phenotype of animal *i* as the giver within group 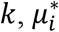 is the expected mean associated with *i*, 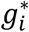 is the additive genetic effect of animal *i* on giving interaction, 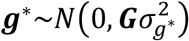 is the random effect of the social group *k*, 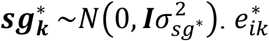 is the error, 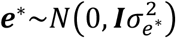. The total variance of *y*_*i*⋅*k*_ is:

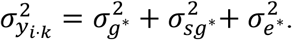

The heritability for total giving interaction (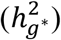) in the giver marginal model is:

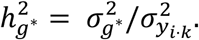

However, a model for *y*_*i*⋅*k*_ can also be derived by summing the terms in Equation (1).

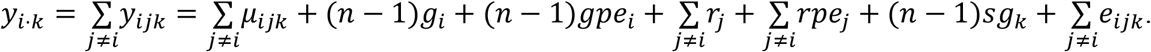

Thereby, the phenotypic variance of *y*_*i*⋅*k*_ under Equation (2) is calculated as:

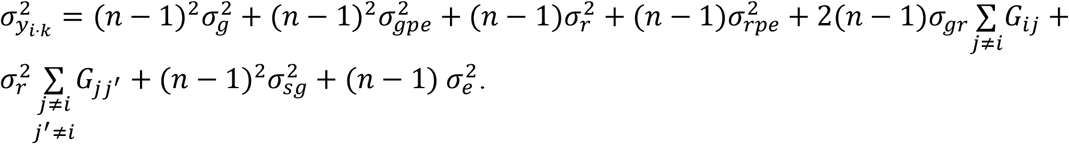

This derivation assumes that group members are unrelated in the base population, such that *G*_*ij*_ and *G*_*jj′*_ are 0. Drawing an equivalence between models (1) and (2) is only possible by assuming the receiver genetic variance, 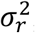, and permanent environmental variance, 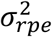, are equal to 0, then, *y*_*i*⋅*k*_ and 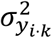 can be written as:

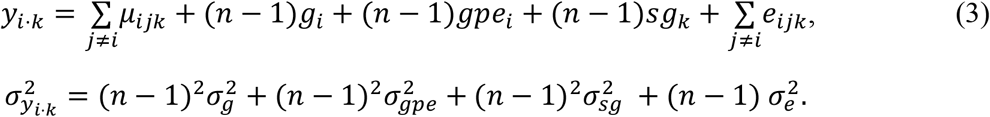

Furthermore, if the permanent environmental effect of the giver is also dismissed 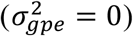, Equation (2) and (3) are equivalent, i.e. 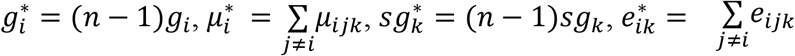. Consequently, the variance components will satisfy: 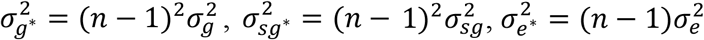. Under these conditions, Equation (3) is a linear reparameterization of Equation (2), and the variance components are rescaled according to group size *n*.

However, when 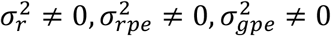, the equivalence will not hold. Moreover, in practical parameter estimation, these components will be omitted from the marginal model, thus, their aggregated explained variation may be redistributed to the variance components retained in the marginal model. As a result, ignoring these components may lead to biased estimates of marginal variance components. The extent of redistribution depends on the variance structures of the omitted components and their collinearity with the retained variance structures. We discuss this further in the Results section and provide a matrix derivation in Appendix Section 2.

#### Column-Sum receiver marginal model

Similarly, the receiver marginal model is obtained by summing the dyadic interaction matrix ***Y***_***k***_ over columns:

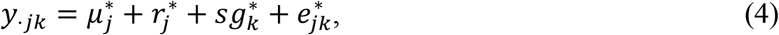

where *y*_⋅*jk*_ denotes the aggregated social interaction phenotype of animal *j* as the receiver within group *k*, 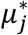 is the overall intercept and fixed effects associated with *j*, 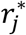 is the additive genetic effect of animal *j* on receiving interaction, 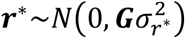 is the random effect of the social group *k*, 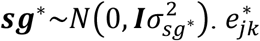 is the error, 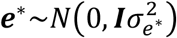. The total variance of *y*_⋅*jk*_ is:

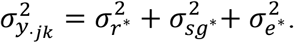

The heritability for total receiving interaction 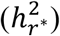 in the receiver marginal model is:

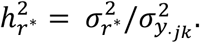

*y*_⋅*jk*_ and the total variance of *y*_⋅*jk*_ are derived based on Equation (1):

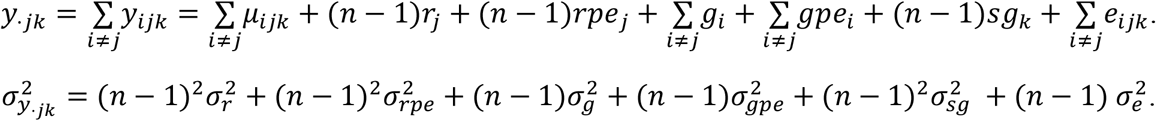

There is published consensus that the giver variance is much larger than the receiver variance (Løvendahl et al. 2005; Turner et al. 2009; Han et al. 2022), thus, we do not anticipate that it will ever be appropriate to assume that the giver genetic variance, 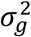, and the giver permanent environmental variance, 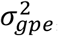, are equal to 0. Therefore, in most practical applications, the receiver marginal model may lead to greater bias in the estimation of genetic variance components, as substantial variance from giver effects is not explicitly accounted for in the model structure.

### Dyadic model for undirected social interactions

The modeling of undirected dyadic data has received less attention in the quantitative genetic literature. However, several sensing technologies are generating data on this type of interactions (Vander Wal et al. 2016; Agha et al. 2025; Perinot et al. 2025). In this section, we assume a dyadic trait with no direction. Each animal has a single genetic effect and a single permanent environmental effect involved in the interactions. Therefore, it is important to incorporate these traits within a genetic framework, and the model is defined as follows.

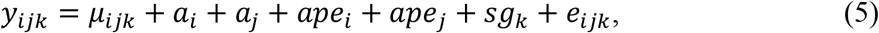

where *y*_*ijk*_ *= y*_*jik*_ is the undirected social interaction phenotype between animal *i* and animal *j* within the *k*th group. *a*_*i*_ and *a*_*j*_ are the random additive genetic effects of the animal *i* and the animal *j*, 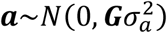. *ape*_*i*_ and *ape*_*j*_ are the random permanent environmental effects of the animal *i* and the animal *j* respectively, with 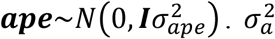 and 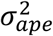 are the animal genetic variance and permanent environmental variance, respectively. The total variance of *y*_*ijk*_ is:

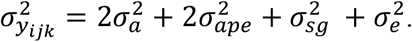

The heritability and repeatability for reciprocal aggression are defined as:

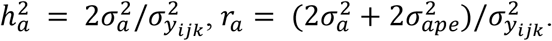

The marginal model for undirected social interaction will be written as:

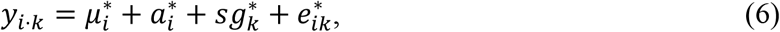

where 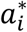 is the additive genetic effect of animal *i*, 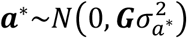. The total variance of *y*_*i*⋅*k*_ given by this marginal model is:

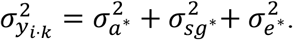

The heritability for total reciprocal interactions in this marginal model is:

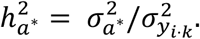

Following Equation (5), *y*_*i*⋅*k*_ can also be derived as:

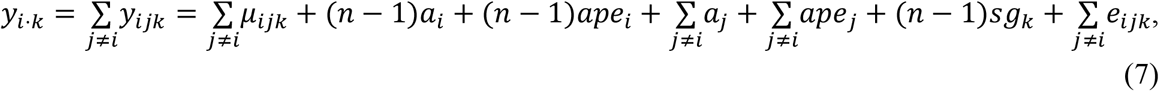

In the base population, we have *cov*(*a*_*i*_, *a*_*j*_) = 0, so the variance of *y*_*i*⋅*k*_ is given by:

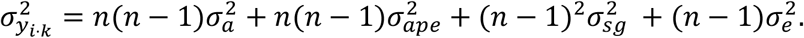

The conditions for the equivalence of Equation (6) and Equation (7) are:

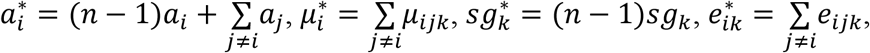

If permanent environmental effect 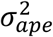 is 0, the variance components satisfy:

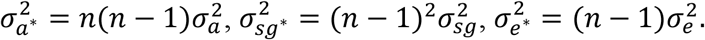

The aggregated animal genetic variance is scaled by *n*(*n* − 1). However, as discussed in the previous section, when 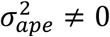, the presented equivalence will not hold. Moreover, the omission of permanent environmental effects and the aggregation of dyadic traits may cause variance components derived from the dyadic model to be redistributed among the variance components retained in the fitted marginal model. We discuss this further in the Results section and provide a matrix derivation in Appendix Section 3.

### Modeling of fixed effects

Fixed effects in dyadic trait models can differ from those in individual-traits models. One key difference is the potential inclusion of dyadic-level covariates. For example, prior affiliation between two individuals can be included as an important dyadic-level covariate, such as whether they previously shared the same social environment. Individual-level covariates can also be incorporated, such as sex, body weight, age, etc. In directed dyadic models, because individuals play distinct roles, the fixed effect coefficients for an animal acting as a giver may differ from those when it acts as a receiver. Therefore, fixed effects for giver and receiver need to be specified separately. However, for undirected interactions, where the roles of the two individuals cannot be distinguished, a single covariate can be used to represent their relative relationship, such as a weight difference or an indicator variable for whether the two individuals are of the same sex. Although these fixed effects are not of direct genetic interest, accounting for them is necessary to obtain unbiased estimates of genetic parameters.

All in all, a direct equivalence between dyadic and marginal models is only possible in very specific cases when some key variance components of the dyadic model are zero and there are no dyadic-level fixed effects. Thus, aggregation of dyadic records into marginal traits should be avoided whenever possible. We illustrate this with an application to a real dataset in the next section.

## Application

In this section we show an application of dyadic data modeling, using data from group-housed pigs that were behaviorally phenotyped. We compare the inferences obtained from dyadic models to those obtained with marginal models.

### Materials and Methods

#### Data

The data used in this study were obtained from previously published studies (Angarita et al. 2019; Han et al. 2022). Video recordings were collected for 9 hours immediately after mixing 797 purebred Yorkshire pigs (406 gilts, 386 barrows) into 59 single-sex finishing social groups. Each group consisted of 10 to 15 pigs (either barrows or gilts) originating from 4 to 6 different nursery groups. Aggressive behaviors were manually annotated from video recordings by 21 trained observers as described by O’Malley et al. (2022). In directed aggression behaviors (e.g., attacks and bites), both the aggressor (giver) and the recipient (receiver) were explicitly identified. For each ordered pair of animals, the aggression duration *y*_*ijk*_ (in seconds) was computed by summing the durations of all annotated directed aggression behaviors during the 9-hour post-mixing period. In undirected aggression behaviors, animals participating in reciprocal fights were identified, with no distinction between the giver and the receiver. The aggression duration *y*_*ijk*_ = *y*_*jik*_ (in seconds) was computed by summing the durations of all annotated undirected aggression behaviors during the 9-hour post-mixing period. In total, the final dataset included 10,032 directed dyadic aggression records and 5,016 undirected dyadic aggression records. The genomic relationship matrix was available from Angarita et al. (2019). Animals were genotyped using the GeneSeek Genomic Profiler for Porcine HD BeadChip with 49,461 SNP. The genomic relationship matrix ***G*** was computed after VanRaden (VanRaden 2008). Additional details were provided in Angarita et al. (2019).

#### Variance component estimation

Dyadic and marginal models were fitted using restricted maximum likelihood methodology in ASREML 4.2 (Gilmour et al., 2015). Fixed effects for directed dyadic model included individual-level covariates: sex, weight of the giver and receiver relative to the average social group weight, and dyadic-level covariates: an indicator of previous litter mates (1 if the animals were litter mates and 0 if they did not share the same litter) and an indicator of previous nursery pen mates (1 if the animals had been nursery pen mates and 0 if they did not share the same nursery group). The undirected dyadic model included the same covariates, except that the weight covariate was defined as the absolute difference between the weights of the two animals. Fixed effects for marginal models included only individuals’ sex and weight. Random effects and variance components in both dyadic and marginal models were as described in equations (1), (2), (4) for directed traits and (5), (6) for undirected traits.

## Results and Discussions

### Directed aggression

Among fixed effects, prior nursery mate was the only significant covariate (negative value, P < 0.001). This means that pigs that had shared a nursery social group showed significantly shorter fight durations after being remixed into grow-finishing groups. Nursery mate is a type of fixed effects that is only available through dyadic modeling, because it consists of an indicator variable taking the value 1 if the two animals in the dyad were housed together in the nursery pen and a 0 otherwise. Thus, it represented an effect of prior acquaintance. Other covariates (weight, sex) were not significant.

In the directed dyadic model (Table 1), the giver genetic effect explained a small but heritable proportion of the phenotypic variation (5.6%), whereas the receiver genetic effect was close to zero, indicating that delivering aggression was more heritable than receiving aggression. Genetic and permanent environmental components together explained 24% of the total variation in giving aggression, with permanent environmental components accounting for more variance than genetic components. The large, persistent environmental variance reflects the influence of early-life experiences on the expression of aggression. In contrast, the permanent environmental variance for receiving aggression was small, leading to low receiver repeatability. The genetic correlation between the giver and receiver was -0.28 but not significant, whereas the permanent environmental correlation between the giver and receiver was 0.31 and significant. This suggests that non-genetic factors influencing an individual’s tendency to engage in longer aggression may also increase the duration of aggression they receive, consistent with previous findings (Han et al. 2022).

**Table 1.** Estimated statistics for (co)variance components explained on dyadic duration of aggression with directed dyadic model (SE in parentheses).

| Parameter |  | Estimates |
| --- | --- | --- |
| $\sigma_g^2$ | Giver genetic variance | 0.079 (0.025) |
| $\sigma_r^2$ | Receiver genetic variance | 0.013 (0.006) |
| $\sigma_{gr}$ | Genetic covariance | -0.009 (0.009) |
| $\sigma_{gpe}^2$ | Giver permanent environmental variance | 0.259 (0.024) |
| $\sigma_{rpe}^2$ | Receiver permanent environmental variance | 0.026 (0.007) |
| $\sigma_{grpe}$ | Permanent environmental covariance | 0.026 (0.009) |
| $\sigma_{sg}^2$ | Social group variance | 0.033 (0.014) |
| $\sigma_e^2$ | Error variance | 0.992 (0.015) |
| $\rho_{gr}$ | Genetic correlation | -0.280 (0.279) |
| $\rho_{grpe}$ | Permanent environmental correlation | 0.312 (0.114) |
| $h_g^2$ | Giver heritability | 0.056 (0.018) |
| $r_g$ | Giver repeatability | 0.241 (0.014) |
| $h_r^2$ | Receiver heritability | 0.009 (0.005) |
| $r_r$ | Receiver repeatability | 0.028 (0.005) |

Estimates of variance components using the giver and receiver marginal models are presented in Table 2 and Table 3, respectively, to illustrate how aggregating records of the social interaction matrix complicates the dissection of genetic and various environmental effects. The estimated variance components from each marginal model were compared with the expected variance components derived from the dyadic model, assuming particular cases of variance component values as described below, and using the average group size in this study (n = 13.5).

**Table 2.** Estimated statistics and expected statistics under assumptions for variance components explained on individuals’ total duration of giving aggression with marginal model (SE in parentheses).

| Parameter | | Estimates | Expected<br>(under $\sigma_r^2 =$<br>$\sigma_{rpe}^2 = \sigma_{gpe}^2 =$<br>0) |
| --- | --- | --- | --- |
| $\sigma_g^{2*}$ | Giver genetic variance | 11.226 (3.755) | 12.343 |
| $\sigma_{sg}^{2*}$ | Social group variance | 6.894 (2.253) | 5.156 |
| $\sigma_e^{2*}$ | Error variance | 53.189 (3.762) | 12.400 |
| $h_g^{2*}$ | Giver heritability | 0.157 (0.050) | 0.413 |

**Table 3.** Estimated statistics and expected statistics under assumptions for variance components explained on individuals’ total duration of receiving aggression with marginal model (SE in parentheses).

| Parameter | | Estimates | Expected<br>(under $\sigma_g^2 =$<br>$\sigma_{rpe}^2 = \sigma_{gpe}^2 =$<br>0) |
| --- | --- | --- | --- |
| $\sigma_r^{2*}$ | Receiver genetic variance | 2.433 (1.058) | 2.031 |
| $\sigma_{sg}^{2*}$ | Social group variance | 11.011 (2.358) | 5.156 |
| $\sigma_{e^*}^2$ | Error variance | 16.400 (1.145) | 12.400 |
| $h_{r^*}^2$ | Receiver heritability | 0.082 (0.035) | 0.104 |

In the Giver marginal model (Table 2), the expected variance components were derived assuming 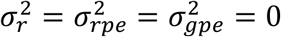. The estimated giver genetic variance in the marginal model 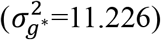 was close to the expected giver genetic variance derived from components estimated in the dyadic model 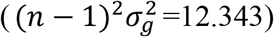. Moreover, both the estimated social group variance and error variance in the marginal model were larger than their expected values. We attributed this to omitting permanent environmental effects from the dyadic model when conducting the marginal analysis. Using arguments explained in the Appendix Section 2.2, the error variance seems to have absorbed the giver permanent environmental variance 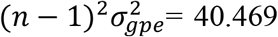. The social group variance apparently absorbed the receiver permanent environmental variance with expected contributions 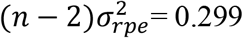 and may have absorbed parts of the aggregated receiver genetic variance. Thereby, under conditions where the assumption that 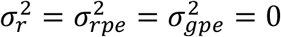 is not plausible, as anticipated from Table 1, the remaining components of the marginal model are explaining other sources of variation in the dyadic data.

In the receiver marginal model (Table 3), the expected variance components were derived under 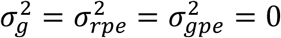. The receiver genetic variance was slightly overestimated, possibly due to the absorption of 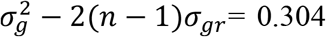, as derived in the Appendix Section 2.2. Both the estimated error variance and estimated social group variance were largely inflated by not accounting for the giver’s and receiver’s permanent environmental effects. The social group variance absorbed the giver permanent environmental variance 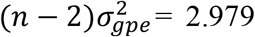 and 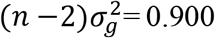, and the error variance absorbed the receiver permanent environmental variance 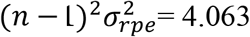.

It was observed that the heritability estimated from the dyadic model was lower than that from the marginal model, which was consistent with results from a previous study (Løvendahl et al. 2005). This is because each model estimates the heritability of a different trait: the dyadic model estimates the heritability of the average duration of single aggression events delivered or received, whereas the marginal model estimates the heritability of the total duration of all aggression events delivered or received by an animal. The error variance in the marginal model is expected to increase by a factor of (*n* − 1), whereas the other variance components increase by a factor of (*n* − 1)^2^ . Therefore, even when the genetic variance from the marginal model is an exact scaled version of that from the dyadic model, the marginal model will still have a larger heritability. In practice, the estimated heritability will also depend on how omitted variance components are redistributed among the fitted variance components. Thus, we suggest that heritability should be interpreted based on estimates from the dyadic model rather than the marginal model.

### Undirected aggression

Fixed effects of shared nursery pen were also negative and significant for undirected aggression (P < 0.001). It means that pigs sharing the same nursery pen had a shorter duration of reciprocal fights once re-grouped with other pigs into finisher pens. A similar test was not available from the marginal models. In the marginal model analysis, the individual’s body weight was significant (P < 0.001), with heavier pigs showing more reciprocal aggression. This reflects that heavier individuals have a greater overall tendency to engage in reciprocal interactions. In contrast, in the dyadic model, absolute weight difference was not significant, suggesting that differences in body weight between two pigs do not strongly determine the duration of mutual interactions.

In the undirected dyadic model (Table 4), the heritability of reciprocal aggression was 0.106, while the repeatability was 0.416, indicating that permanent environmental effects also played an important role in the undirected aggression.

**Table 4.** Estimated statistics for variance components explained on dyadic reciprocal aggression with the undirected dyadic model(SE in parentheses).

| Parameter |  | Estimates |
| --- | --- | --- |
| $\sigma_a^2$ | Animal genetic variance | 0.234 (0.069) |
| $\sigma_{ape}^2$ | Animal permanent environmental variance | 0.682 (0.065) |
| $\sigma_{sg}^2$ | Social group variance | 0.175 (0.092) |
| $\sigma_e^2$ | Error variance | 2.400 (0.058) |
| $h_a^2$ | Animal heritability | 0.106 (0.030) |
| $r_a$ | Animal repeatability | 0.416 (0.021) |

In the marginal model (Table 5), the estimated animal genetic variance was not equal to the expected value derived from the dyadic model under the assumption that the animal permanent environmental variance is 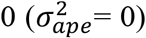. In the theoretical background section, we derived that for a focal animal in the base population, and assuming that permanent-environmental variance is zero, we expect the marginal animal genetic variance to be 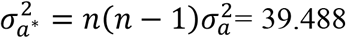. However, this result does not necessarily hold for an arbitrary set of related animals. As shown in Appendix 3.1, only 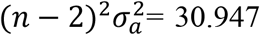 can be directly redistributed to 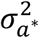. In addition, because the marginal model did not account for animal-specific permanent environmental effects, both the social group variance and the error variance were overestimated. The theoretical permanent environmental variance 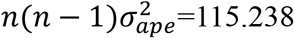 was redistributed to the social group variance and error variance.

**Table 5.** Estimated statistics and expected statistics under assumptions for variance components explained on individuals’ total duration of reciprocal aggression with marginal model(SE in parentheses).

| Parameter | | Estimates | Expected<br>(Under<br>$\sigma_{ape}^2=0$) |
| --- | --- | --- | --- |
| $\sigma_{a^*}^2$ | Animal genetic variance | 33.643 (10.134) | 39.488 |
| $\sigma_{sg^*}^2$ | Social group variance | 61.510 (13.916) | 27.344 |
| $\sigma_{e^*}^2$ | Error variance | 115.883 (8.860) | 30.000 |
| $h_{a^*}^2$ | Animal heritability | 0.159 (0.046) | 0.408 |

Overall, based on the application’s estimated results and the discussions in Appendix Sections 1, 2, and 3, we can summarize several findings. When aggregating directed interactions for a given role, the dyadic genetic variance corresponding to that focal role can be directly retained in the genetic variance of the marginal model, and the dyadic social group and error variances can also be retained in their corresponding marginal variance components. However, the omitted variance components will be redistributed to varying degrees among the retained variance components in the marginal model. When aggregating undirected interactions, all dyadic variance components will be redistributed among the variance components retained in the marginal model. Although we demonstrated in the Appendix that some variance parameters between the dyadic and marginal models have predictable linear (scalar) or non-linear relationships, these approximations also depend on the specific variance structures of the data. Therefore, when dyadic records are available, the dyadic model should be preferred over the marginal model to avoid confounding issues caused by aggregation.

We also compared breeding values estimated from the dyadic and marginal model (see Section 4 Figure 1 in the Appendix). For this exemplary dataset, the correlations between the estimated 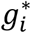 and *g*_*i*_, between *r*^⋅^ and *r*_*i*_, and between 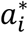 and *a*_*i*_ was 0.98, 0.98, 0.97. We attribute this to the fact that in this example, the estimated marginal giver and receiver variances did not absorb a significant portion of other components. However, with a different data structure (e.g., a different distribution of within- and between-group relations), the results can change, and we recommend always using the dyadic model when dyadic-level data are available.

**Figure 1.**
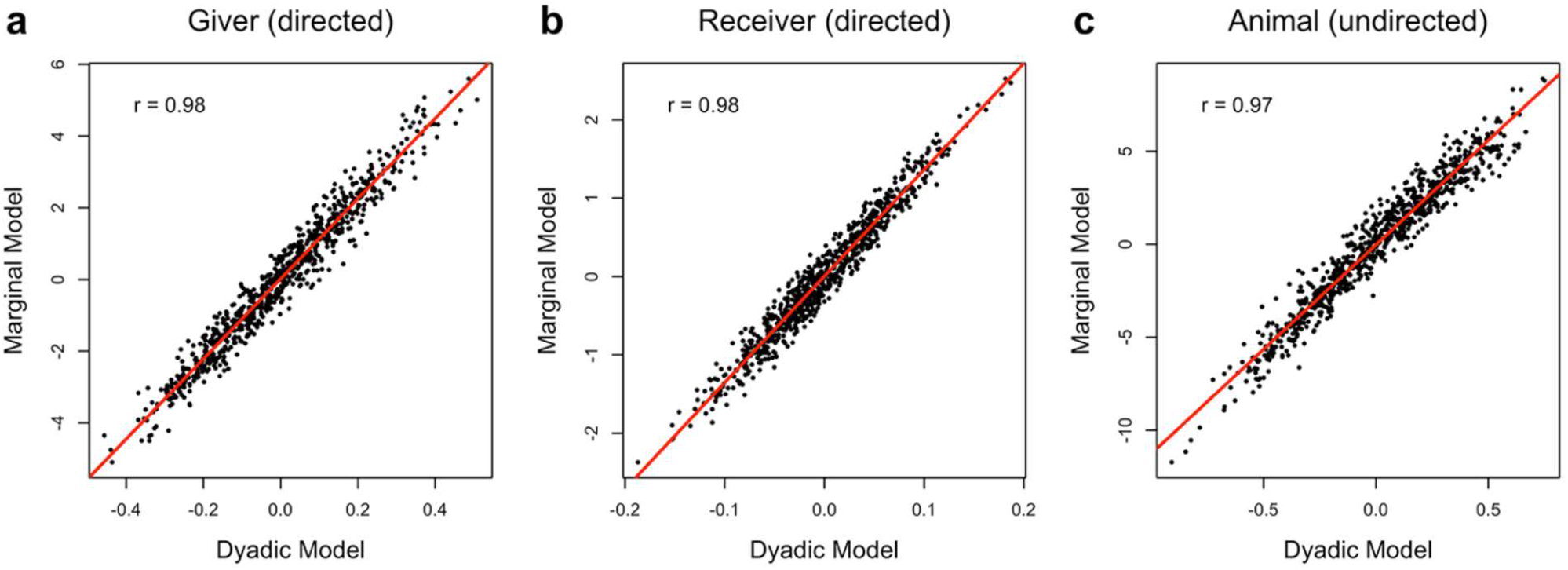
Comparison of BLUP for genetic effects obtained from dyadic models and marginal models.

## Conclusion

This study presents the dyadic model for both directed and undirected social interaction traits and demonstrates why the dyadic model is a more appropriate modeling strategy than the more commonly used marginal model based on aggregated interactions. The dyadic model provides the clearest and most interpretable variance decomposition, allowing the estimation of genetic effects across different roles and permanent environmental effects, which can improve our understanding of the genetic and non-genetic mechanisms underlying social interactions. In contrast, aggregation of dyadic interaction traits may redistribute variance components, posing challenges for interpreting the true sources of variance. This framework can be extended to other pairwise phenotypes, such as maternal–progeny traits and disease transmission traits, which will be helpful for breeding decisions and farm management.

## Data availability statement

All scripts and data for implementing the models are available at: https://github.com/xiaohanj-isu/social_interaction.

## Author contributions

XHJ performed the analyses and drafted the manuscript. JPS conceived the study and supervised the research. JMS contributed to the original experimental design and data collection. XHJ, JMS and JPS contributed to the interpretation of the results. All authors read and approved the final manuscript.

## Appendices

## 1. Estimation of variance parameter and confounding assessment using Average Information REML

The directed dyadic social interaction model can be written in matrix format:

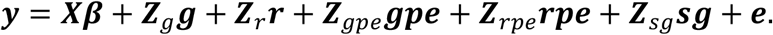

where ***y*** is the vector of directed dyadic social interaction records. ***β*** is the vector of fixed effects, ***X*** represents the incidence matrix relating the fixed effects to each individual. ***g*** and ***r*** are the vectors of additive genetic effects for the giver and receiver, ***Z***_*g*_ and ***Z***_*r*_ are the incidence matrixes relating records in ***y*** to the additive genetic effects ***g*** and ***r. gpe*** and ***rpe*** are the vectors of permanent environmental effects for the giver and receiver, ***Z***_*gpe*_ and ***Z***_*rpe*_ are the incidence matrixes relating records in ***y*** to the additive permanent environmental effects ***gpe*** and ***rpe. sg*** is the vector of social group effects, ***Z***_*sg*_ is the incidence matrix relating records in ***y*** to the social group effects. ***e*** is the vector of error. The same incidence structure is used for the giver’s genetic and giver’s permanent environmental effects, ***Z***_*g*_ *=* ***Z***_*gpe*_, and similarly for the receiver, ***Z***_*r*_ *=* ***Z***_*rpe*_.

Assuming:

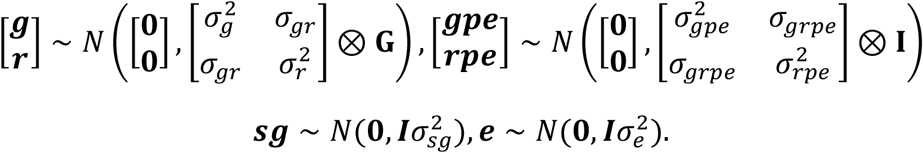

The vector of (co)variance parameters is

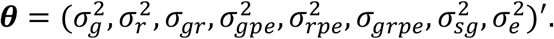

The variance matrix of ***y*** is

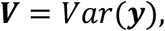

which can be written as

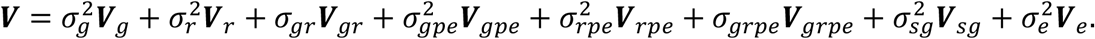

where

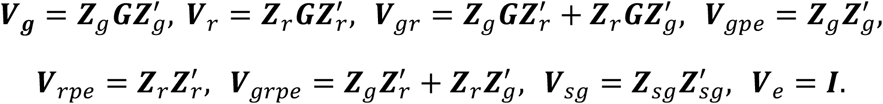

Here we briefly review the AI-REML algorithm in Gilmour et al. (1995), as it provides the foundation for the subsequent explanations. The log residual likelihood (ignoring constants) can be written as

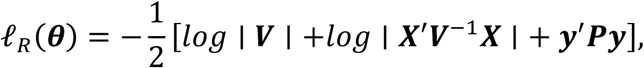

where

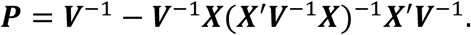

The first derivative of *ℓ*_*R*_(***θ***) gives the score *U*_*i*_(***θ***)

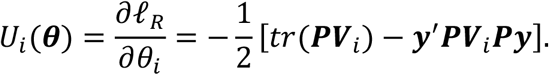

where

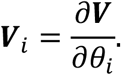

Thus, if 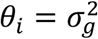, then 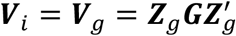. If 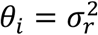, then 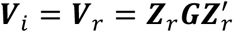. The REML estimates of *θ*_*i*_ satisfy *U*_*i*_(***θ***) = 0.

In the Average Information REML (AI-REML) algorithm, the average information matrix ***I***_***A***_ was calculated to update ***θ***. The elements of ***I***_***A***_ are given by

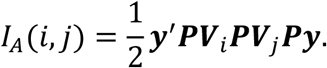

The AI-REML update of ***θ*** can be written as

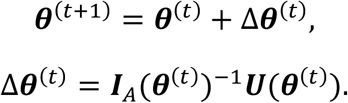

Thus, the REML estimate of each variance parameter *θ*_*i*_ depends on its own variance structures ***V***_*i*_, but also on the full variance matrix ***V***, the fixed effects matrix ***X*** and records vector ***y***. As a result, the estimation of different variance parameters can influence each other through their contributions to the overall variance structure. In general, the more similar two ***V***_*i*_ matrices are, the more difficult it is for REML to disentangle the corresponding variance parameter, as similar variance structures make the average information matrix nearly singular. For example, the difference between 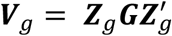 and 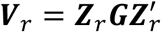 comes from their incidence matrices ***Z*** and ***Z***_*r*_ . Similarly, the difference between 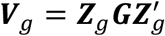 and 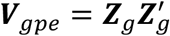 is determined by the genomic relationship matrix ***G***. If genomic relationships between individuals are close to 0, ***G*** becomes close to an identify matrix, making it difficult to uniquely identify genetic variance from permanent environmental variance. Moreover, the separating 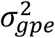 and 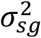 is influenced on and ***Z***_*sg*_, as 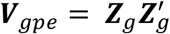 and 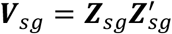, and the same applies to the separation of 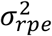 and 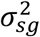 through ***Z***_*r*_ and ***Z***_*sg*_. Therefore, distinguishing these two parameters requires repeated records for multiple individuals within each social group.

This collinearity can be assessed through the sampling correlations among the estimated variance parameters derived from the inverse of the average information matrix. When the number of records is large, we have the approximation 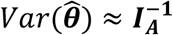. Thus, the inverse of ***I***_***A***_ provides an approximate variance-covariance matrix of estimated variance parameters and can be used to calculate their correlations. High absolute correlations indicate that parameters are highly confounded, meaning the model has limited ability to disentangle their effects (Lee et al. 2010).

## 2. Expected redistribution of dyadic variance components in the fitted marginal model

This derivation is presented for the giver marginal model, assuming *m* social groups of equal size *n*, where each animal interacts with all other (*n* − 1) animals within its group. The total number of social groups is *m*. Because the receiver marginal model is obtained by exchanging the roles of giver and receiver, all derivations apply analogously to the receiver marginal model. For the giver marginal model, each marginal record is obtained by summing all dyadic records in which the animal acts as the giver:

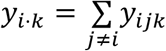

Let ***M*** be the aggregation matrix that maps dyadic records to giver marginal records, so the giver marginal record vector ***y***^⋅^ = ***My*** . Each row of ***M*** corresponds to a giver marginal record (*i* ⋅ *k*), and each column corresponds to a dyadic record (*ijk*), so 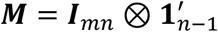, where ***I***_*mn*_ is the identity matrix of order *m* × *n, m* is the number of groups, and *n* is the number of animals per group. Multiplying the dyadic model by ***M***, we obtain

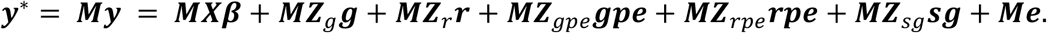

The variance matrix of the giver marginal record derived from the dyadic model is

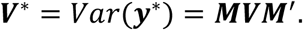

which can be written as

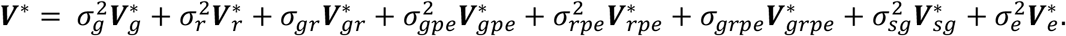

where

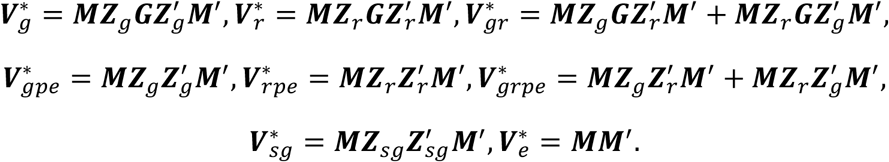

The giver marginal model we indeed fit is

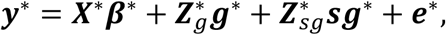

with assuming

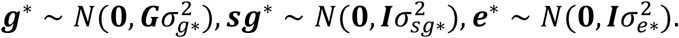

In this fitted giver marginal model, we have 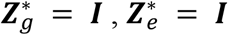, then 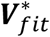 is

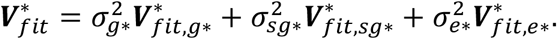

where

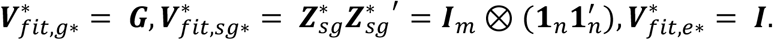

Therefore, the fitted marginal model can be viewed as a reduced dyadic model, with some variance components are omitted and redistributed among the retained variance components in the fitted marginal model. The potential redistribution of omitted variance components can be understood by comparing the variance structures derived by the dyadic model with those obtained in the fitted marginal model.

### 2.1 Redistribution of 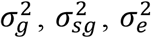

For the giver genetic variance, 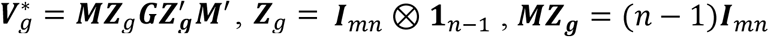. Therefore, 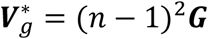. This variance structure is proportional to the fitted marginal genetic covariance structure 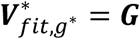. The fitted marginal giver genetic variance is 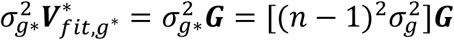. This indicates that the fitted marginal genetic variance 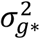 will absorb 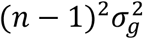. Similarly, for the social group variance, 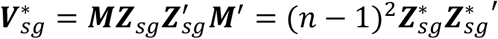, and 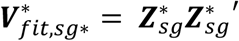, showing that the fitted marginal social group variance 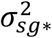 will absorb 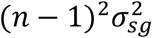. For the error variance, 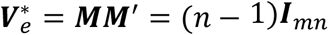, indicating that the fitted marginal error variance 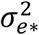 will absorb 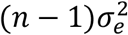.

For the variance components omitted from the fitted giver marginal model, including 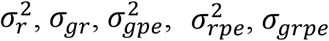, we follow the same logic to examine the collinearity between their variance structures and the variance structures retained in the fitted marginal model.

### 2.2 Redistribution of 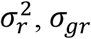

The receiver genetic variance is 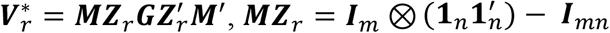, so 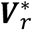 can be derived as

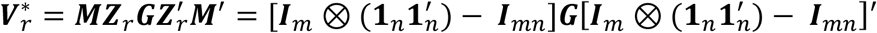

We can also derive the fitted marginal social group variance structure 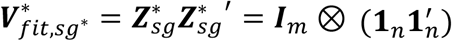. Then 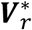 can be reorganized as

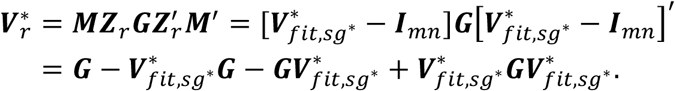

In this structure, only ***G*** is exactly proportional to the fitted marginal genetic variance structure 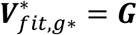. Thus, 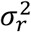 can be directly redistributed to 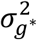. In the base population, when ***G*** = ***I***, the remaining terms will be 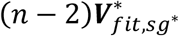. Therefore, 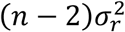 will be absorbed by the fitted social group variance 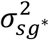. However, when ***G*** ≠ ***I***, these remaining terms involve products between ***G*** and the fitted social group structure 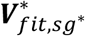, so they may be absorbed by 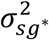, or partially by 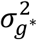, depending on the collinearity among the fitted variance structures.

Similarity, the giver and receiver genetic covariance structure is

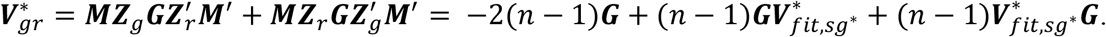

Therefore, −2(*n* − 1)*σ*_*gr*_ will be redistributed to the fitted social group variance 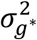. The remaining variance may be absorbed by 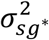, or partially by 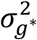.

### 2.3 Redistribution of 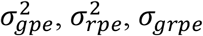

For the giver permanent environmental variance, 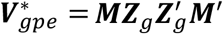. Since ***MZ***_*g*_ = (*n* − 1)***I***_*mn*_, we have 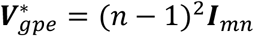. This structure is proportional to the fitted marginal error variance structure 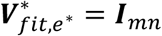. Therefore, the fitted marginal error variance 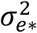 can absorb 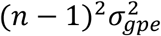.

The receiver permanent environmental variance structure 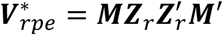 can be rewritten as 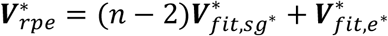, indicating that the fitted marginal social group variance 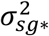 may absorb 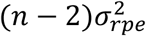, and the fitted marginal error variance 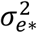 may absorb 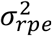. Similarly, the giver-receiver permanent environmental covariance structure, 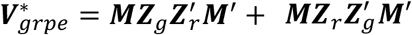, can be rewritten as 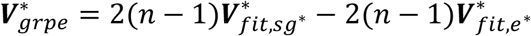, indicating that the fitted marginal social group variance 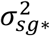 may absorb 2(*n* − 1)*σ*_*grpe*_, and the fitted marginal error variance 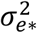 may absorb −2(*n* − 1)*σ*_*grpe*_.

In summary, the fitted marginal genetic variance 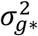 should contain 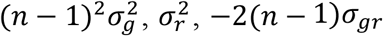, and may contain parts of the remaining aggregated receiver genetic variance and giver-receiver genetic covariance. The fitted marginal social group variance 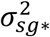 should contain 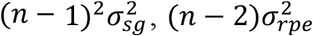, and 2(*n* − 1)*σ*_*grpe*_, and may contain parts of the remaining aggregated receiver genetic variance and giver-receiver genetic covariance. The fitted marginal error variance 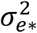 should contain 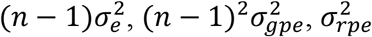, and −2(*n* − 1)*σ*_*grpe*_.

It should be noted that the fitted marginal model here is based on the aggregation of the dyadic model. In practice, the fixed effects ***X***^⋅^***β***^⋅^ of the fitted marginal model is not completely equivalent to the aggregated fixed effects matrix ***Xβ*** from the dyadic model, as the dyadic model can include dyadic level covariates. Therefore, the REML estimates from the fitted marginal model may deviate from the theoretical expectations. In addition, the derivations above assume balanced group size. When group size varies across groups, the exact scalar relationships will not hold. Nevertheless, these algebraic relationships based on variance structures still provide useful insight into the expected directions of redistribution and the potential sources of confounding among variance components.

## 3. Expected redistribution of undirected dyadic variance components in the fitted animal marginal model

The undirected dyadic social interaction model can be written in matrix format:

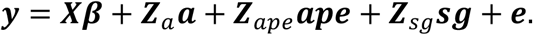

where ***y*** is the vector of undirected social interaction records. ***a*** is the vector of additive genetic effects, ***Z***_*a*_ is the animal incidence matrix relating undirected dyadic records in ***y*** to the animal genetic effects. Each row of ***Z***_*a*_ corresponds to one dyadic record, with entries equal to 1 for animals *i* and *j* . ***ape*** is the vector of animal permanent environmental effects, ***Z***_*ape*_ is the incidence matrixes relating records in ***y*** to the animal permanent environmental effects, with ***Z***_*a*_ = ***Z***_*ape*_. ***sg*** is the vector of social group effects, ***Z***_*sg*_ is the incidence matrix relating records in ***y*** to the social group effects. ***e*** is the vector of error.

Assuming:

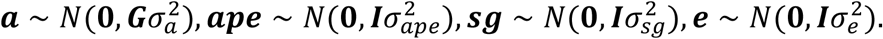

The variance matrix of the undirected dyadic record is

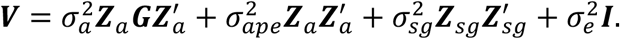

Let ***M*** be the aggregation matrix that maps undirected dyadic records to marginal records, so that

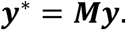

Multiplying the dyadic model by ***M***, we obtain

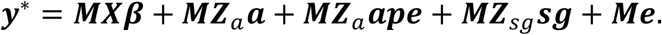

Thus, the variance matrix of the marginal records is

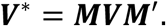

which can be written as

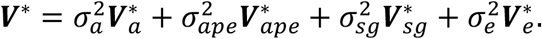

where

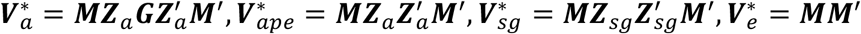

For balanced group size *n* with *m* social groups,

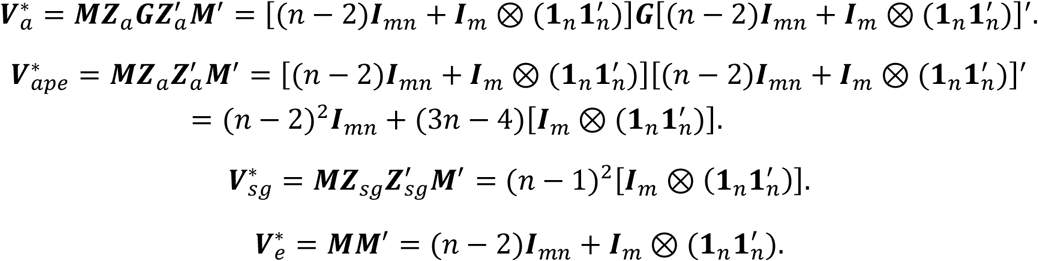

The fitted marginal model is written as

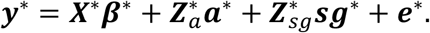

with assuming

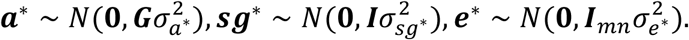

We have 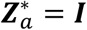, then the variance structure 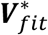 is

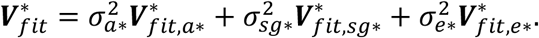

where

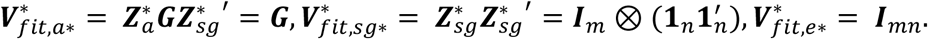

By comparing 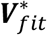 and ***V***^⋅^, we can infer how aggregated variance components from the dyadic model may be redistributed to the fitted marginal variance components.

### 3.1 Redistribution of 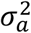

The dyadic genetic variance structure 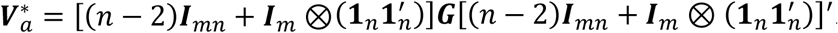.

In the base population, where animals are assumed to be unrelated ***G*** = ***I***, we have

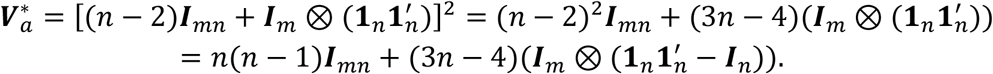

Therefore, only the diagonal elements in 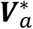 correspond to 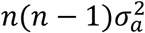, which is the variance of an aggregated record for a single animal, as derived in the main text, we show 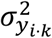:

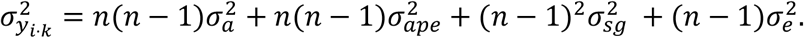

The off-diagonal elements in 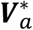 represent covariances between aggregated records of different animals within the same social group.

However, when ***G*** ≠, 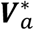 can be rewritten as:

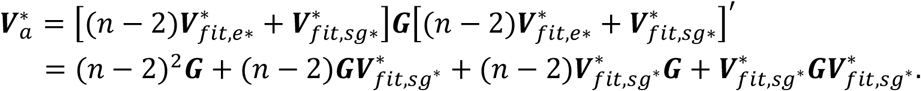

In this structure, only (*n* − 2)^2^***G*** is exactly proportional to the fitted marginal genetic variance structure 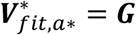. Thus, 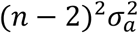 can be directly redistributed to 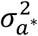. The remaining terms are not simply proportional to ***G***, but involve products between ***G*** and the fitted social group structure 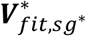. Therefore, these remaining aggregated dyadic genetic variances may be redistributed to 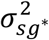, or partially to 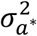, depending on the collinearity among the fitted variance structures (e.g: how animals of certain relationships are assigned to social groups).

### 3.2 Redistribution of 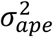

The dyadic permanent environmental variance structure 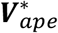 can be rewritten as 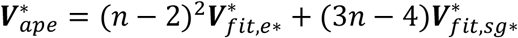, indicating that the fitted error variance 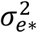 may absorb 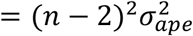, and the fitted marginal social group variance 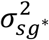 may absorb 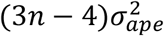. Thus, the total 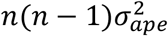 will be redistributed to the fitted social group and error variance.

### 3.3 Redistribution of 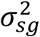

Since 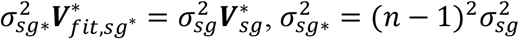, the fitted marginal social group variance 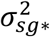 will absorb 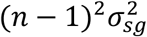.

### 3.4 Redistribution of 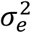

The dyadic error variance structure 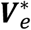 can be rewritten as, 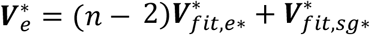, indicating that the fitted marginal error variance 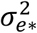 may absorb 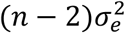, and the fitted marginal social group variance 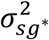 may absorb 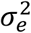.

In summary, the fitted marginal genetic variance 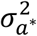 should contain 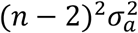, and may also contain parts of the remaining aggregated dyadic genetic variance. The fitted marginal social group variance 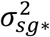 should contain 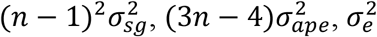, and may also contain parts of the remaining aggregated dyadic genetic variance. The fitted marginal error variance 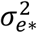 should contain 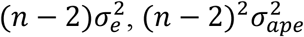.

## References

Agha S et al. 2025. Revealing the Hidden Social Structure of Pigs with AI-Assisted Automated Monitoring Data and Social Network Analysis. Animals. 15(7). 10.3390/ani15070996

Agha S, Fàbrega E, Quintanilla R, Sánchez JP. 2020. Social network analysis of agonistic behaviour and its association with economically important traits in pigs. Animals. 10(11):1–13. 10.3390/ani10112123

Angarita BK et al. 2019. Estimation of indirect social genetic effects for skin lesion count in group-housed pigs by quantifying behavioral interactions. J Anim Sci. 97(9):3658–3668. 10.1093/jas/skz244

Bijma P, Muir WM, Van Arendonk JAM. 2007. Multilevel selection 1: Quantitative genetics of inheritance and response to selection. Genetics. 175(1):277–288. 10.1534/genetics.106.062711

Canario L et al. 2020. Prospects for the Analysis and Reduction of Damaging Behaviour in Group-Housed Livestock, With Application to Pig Breeding. Front Genet. 11. 10.3389/fgene.2020.611073

Canario L, Lundeheim N, Bijma P. 2017. The early-life environment of a pig shapes the phenotypes of its social partners in adulthood. Heredity (Edinb). 118(6):534–541. 10.1038/hdy.2017.3

D’Eath RB, Lawrence AB. 2004. Early life predictors of the development of aggressive behaviour in the domestic pig. Anim Behav. 67(3):501–509. 10.1016/j.anbehav.2003.06.010

Gilmour AR, Thompson R, Cullis BR. 1995. Average Information REML: An Efficient Algorithm for Variance Parameter Estimation in Linear Mixed Models. Biometrics. 51(4):1440. 10.2307/2533274

Han J et al. 2022. Analysis of social interactions in group-housed animals using dyadic linear models. Appl Anim Behav Sci. 256. 10.1016/j.applanim.2022.105747

Hollifield MK et al. 2024. Estimating genetic parameters of digital behavior traits and their relationship with production traits in purebred pigs. Genetics Selection Evolution. 56(1):29. 10.1186/s12711-024-00902-w

Kenny DA, Mohr CD, Levesque MJ. 2001. A social relations variance partitioning of dyadic behavior. Psychol Bull. 127(1):128–141. 10.1037/0033-2909.127.1.128

Kjaer JB, Sørensen P, Su G. 2001. Divergent selection on feather pecking behaviour in laying hens (Gallus gallus domesticus). Appl Anim Behav Sci. 71(3):229–239. 10.1016/S0168-1591(00)00184-2

Lee SH, Goddard ME, Visscher PM, van der Werf JH. 2010. Using the realized relationship matrix to disentangle confounding factors for the estimation of genetic variance components of complex traits. Genetics Selection Evolution. 42(1):22. 10.1186/1297-9686-42-22

Løvendahl P et al. 2005. Aggressive behaviour of sows at mixing and maternal behaviour are heritable and genetically correlated traits. In: Livestock Production Science. Vol. 93. Elsevier; p 73–85. 10.1016/j.livprodsci.2004.11.008

Marina H, Fikse WF, Rönnegård L. 2024. Social network analysis to predict social behavior in dairy cattle. JDS Communications. 5(6):608–612. 10.3168/jdsc.2023-0507

O’Malley CI et al. 2022. The Social Life of Pigs: Changes in Affiliative and Agonistic Behaviors following Mixing. Animals. 12(2). 10.3390/ani12020206

Perinot E, Petelle MB, Gómez Y, Toscano MJ. 2025. Temporal-spatial associations of large groups of laying hens in a quasi-commercial barn. Appl Anim Behav Sci. 283:106516. 10.1016/j.applanim.2025.106516

Turner SP et al. 2009. Genetic validation of postmixing skin injuries in pigs as an indicator of aggressiveness and the relationship with injuries under more stable social conditions. J Anim Sci. 87(10):3076–3082. 10.2527/jas.2008-1558

VanRaden PM. 2008. Efficient methods to compute genomic predictions. J Dairy Sci. 91(11):4414–4423. 10.3168/jds.2007-0980

Vander Wal E, Gagné-Delorme A, Festa-Bianchet M, Pelletier F. 2016. Dyadic associations and individual sociality in bighorn ewes. Behavioral Ecology. 27(2):560–566. 10.1093/beheco/arv193

Wang Z, Doekes H, Bijma P. 2023. Towards genetic improvement of social behaviours in livestock using large-scale sensor data: data simulation and genetic analysis. Genetics Selection Evolution. 55(1). 10.1186/s12711-023-00840-z

Wurtz KE et al. 2017. Estimation of genetic parameters for lesion scores and growth traits in group-housed pigs. J Anim Sci. 95(10):4310–4317. 10.2527/jas2017.1757

